# Petasol Exhibits Rapid and Efficient Epithelial Transport in Epithelial Barrier Models

**DOI:** 10.1101/2025.10.13.682130

**Authors:** Júlia Tárnoki-Zách, Imre Boldizsár, Gábor M Kovács, Szilvia Bősze, András Czirók

## Abstract

The epithelial permeability of petasol, a sesquiterpene derivative with reported antiviral properties, remains largely unexplored despite its potential as a therapeutic candidate. Here, we systematically characterize the transport of petasol across epithelial barriers using a combination of *in vitro* transwell models, automated millifluidic sampling, mathematical modeling, optical- and HPLC-mass spectroscopy. Vero E6 and Caco-2 cell layers were employed as epithelial barrier models, with transepithelial electrical resistance (TEER) measurements confirming the formation of functionally intact epithelial layers. Mathematical modeling of passive diffusion and cellular uptake allowed estimation of key transport parameters, including permeability coefficients and intracellular sequestration rates. Trans-barrier transport of petasol was compared to that of nitazoxanide, chloroquine, and FITC-dextran. Our results reveal that petasol exhibits efficient and rapid transport across both epithelial cell types without substantial intracellular accumulation or metabolic degradation, suggesting favorable properties for systemic delivery.

## 1. Introduction

The COVID-19 pandemic has reinvigorated efforts to discover novel antiviral compounds, a pursuit further intensified by the rising frequency of virus-driven diseases affecting both human and animal populations, including outbreaks of avian influenza and other zoonotic pathogens. Petasol, and its ester petasin, are sesquiterpene derivatives first extracted from Petasites species, has attracted attention due to its neuroprotective, anti-inflammatory, and recently reported antibacterial and antiviral properties [1–3]. Preclinical in vitro studies suggest that petasol may interfere with viral replication pathways, positioning it as a promising candidate for therapeutic development [4]. However, petasol’s pharmacokinetic properties remained unknown, including its ability to cross epithelial barriers – an essential determinant of bioavailability.

For any orally or topically administered drug, transport across epithelial layers is a critical factor that governs systemic absorption and target tissue accessibility. The intestinal epithelium, represented in vitro by cell lines such as Caco-2, forms a selective barrier regulated by tight junctions and active transport processes [5]. Vero E6 cells are frequently used in viral infection studies and – while exhibiting lower TEER values – offer a model for less restrictive epithelial barriers. Measurment and quantitative analysis of a compound’s permeability across in vitro barrier models can reveal key properties such as passive diffusion capacity, cellular uptake, binding, and metabolism [6]. These insights not only inform formulation strategies but also help predict in vivo pharmacokinetics and therapeutic efficacy.

In this study, we use a millifluidic sampling system [7] to monitor the transport of petasol across epithelial cell layers cultured on transwell inserts. For context and comparison, we include three additional well-studied compounds: the clinically used antiviral nitazoxanide [8,9], the chloroquine antiviral with reported intracellular accumulation [10,11], and FITC-dextran – a high-molecular-weight paracellular transport control [12,13]. Using both empirical data and mathematical modeling, we aim to characterize petasol’s epithelial permeability.

## 2. Materials and methods

### 2.1. Reagents, buffers and media

For the *in vitro* assays Dulbecco’s Modified Eagle’s Medium (DMEM) with or without phenol red, phosphate buffered saline (PBS), and L-glutamine were obtaied from Lonza (Basel, Switzerland). Pyruvate and trypsin were obtained from Sigma-Aldrich (St Louis, MO, USA). Non-essential amino acids, fetal bovine serum (FBS) and Penicillin/Streptomycin (10,000 units penicillin and 10 mg streptomycin/mL) were purchased from Gibco (Thermo Fisher Scientific, Waltham, MA, USA). Alamar Blue (rezazurin), Chloroquine, Nitazoxanide, and FITC-dextran were obtained from Sigma-Aldrich (Budapest, Hungary).

### 2.2. Transwell culture

Green monkey (*Chlorocebus sabaeus*) kidney epithelium VERO E6 cells [14–17] were kindly provided by Bernadett Pályi and Zoltán Kis (National Center for Public Health and Pharmacy (NCPHP), Budapest, Hungary), and were obtained from the European Collection of Authenticated Cell Cultures (ECACC 85020206). Caco-2 cells were purchased from Sigma-Aldrich (Budapest, Hungary) as a human epithelial cell line originally derived from a colon carcinoma. In confluent culture, they spontaneously differentiate into a polarized monolayer with features of the absorptive small intestine, including tight junctions and specific enzyme activities.

Vero E6 cells were kept in DMEM high-glucose (4.5 g/L) medium containing 10% FBS supplemented with 100 µg/ml penicillin/streptomycin, 2 mM of L-glutamine. For maintaining Caco-2 cell cultures DMEM high-glucose (4.5 g/L) with phenol red was supplemented with 10% FBS, 2 mM L-glutamine, 100 µg/mL Penicillin/Streptomycin, 1 mM Pyruvate and 1% non-essential amino acids. Media described above, hereafter referred to as complete medium (CM). The same medium without serum and non-essential amino acids is referred to as incomplete medium (ICM). In the transwell insert (both apical and basolateral chambers), the CM was changed every second day in both Vero E6 and Caco-2 cultures. Cultures were kept in humidified, 5% CO2 incubator at 37 °C.

For barrier transport experiments, the polycarbonate transwell inserts (0.6 cm^2^ area, 0.4 µm pore size, Merck Millipore) were equilibrated for at least 2 hours in 12 well plates (Sarstedt, Nümbrecht, Germany) with CM, then into each transwell insert 75 000 cells were seeded in 450 µL CM. At the same time, 1800 µL CM was added to the basolateral compartment. Cultures were grown for 5-7 days under normal culture conditions and were daily monitored by TEER measurements. Immediately before barrier transport measurements, transwell cell cultures and saturated cell free transwell inserts were washed twice with PBS, then the apical and basolateral chambers were filled with 250 µL and 1200 µL ICM, respectively.

For cell-free transport measurements transwell inserts were saturated with serum proteins by incubation in CM for 72 hours. Unsaturated control transwell inserts were incubated in ICM.

### 2.3. Cytotoxicity assay

Cytotoxicity of chloroquine, nitazoxanide, petasol, and FITC–dextran was evaluated using the Alamar Blue (Resazurin) assay on Vero E6 and Caco-2 epithelial cell layers. Cells were seeded in 96-well plates at a density of 5 000 cells per well and cultured at 37 °C for 7 days (Vero E6) or 21 days (Caco-2), CM was replaced every third day. Cells were then treated with compound solutions prepared in ICM at final concentrations ranging from 128 pM to 100 µM in a total volume of 200 µL, ICM alone served as the control. After treatment, the compounds were washed out twice with ICM, and 22.5 µL of Alamar Blue solution (0.15 mg/mL in PBS, pH 7) was added to each well. Following overnight incubation, fluorescence was measured using a Synergy H4 multimode microplate reader (BioTek, Winooski, VT, USA) at excitation/emission wavelengths of 530/30 nm and 610/10 nm. Cytotoxicity (%) was calculated as [1− (*FLUO*_*treated*_/*FLUO*_*control*_)]×100. Four parallel measurements were performed in two independent experiments, and *IC*_50_ values were obtained from the resulting dose–response curves. For the Vero E6 7-day layers, chloroquine exhibited an *IC*_50_ of 48.7 µM, while nitazoxanide and petasol both showed *IC*_50_ > 100 µM; FITC–dextran did not induce measurable cytotoxicity up to 1 mg/mL. In Caco-2 21-day layers, chloroquine and nitazoxanide showed *IC*_50_ values of 58.7 µM and 35.1 µM, respectively, whereas petasol and FITC–dextran again displayed *IC*_50_ > 100 µM and > 1 mg/mL.

### 2.4. Transepithelial electrical resistance

Transepithelial electrical resistance (TEER) was measured using a TEER scanner, a prototype automated scanning system (BioPhys-Concepts Kft, Budapest, Hungary, biophys-concepts.com) equipped with MERSSTX01 chopstick electrodes (Merck, Darmstadt, Germany) connected to a Millicell ERS-2 voltohmmeter (Merck, Darmstadt, Germany). During measurements, the 12-well plate was placed on a heated stage to maintain a constant temperature of 37 °C and transwell inserts were positioned within the wells using special transwell positioners to ensure consistent electrode placement.

As described in detail in [18], each measurement session involved two full scans of both the cell-free (blank) inserts and the inserts containing epithelial barriers, followed by a final third scan of the blank inserts. The entire scanning process for a 12-well plate was completed within approximately 5 minutes. To account for slow temporal drifts in baseline electrical resistance, interpolation was performed based on the resistance values obtained from the blank inserts. TEER values were calculated by subtracting the average resistance of the cell-free inserts from that of the cell-containing insert and normalizing the difference to the surface area of the transwell membrane, 60 mm^2^.

**Figure 1.**
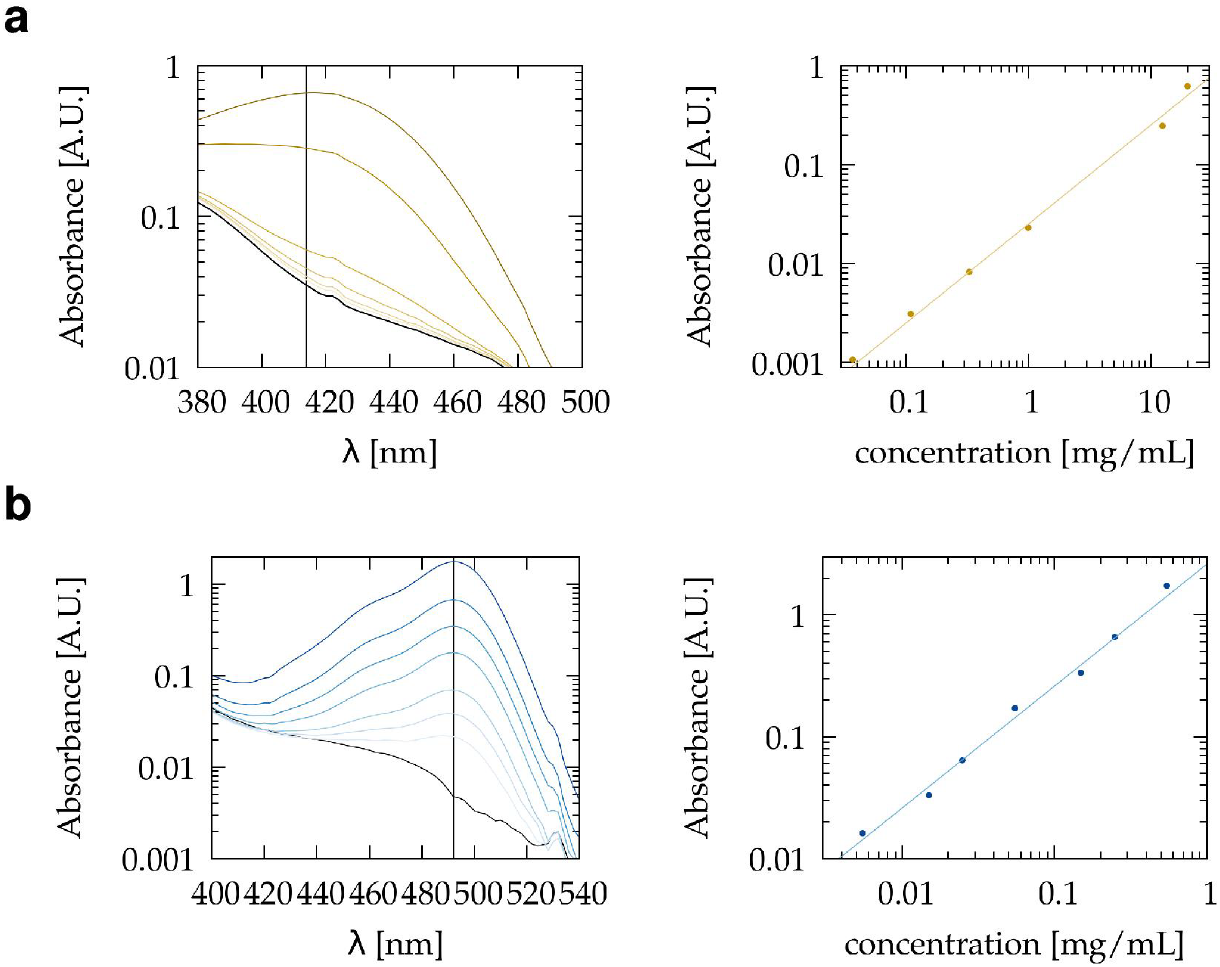
Calibration measurements to establish concentration–absorbance relationships and verify linearity within the range used for transport experiments. Absorbance of the indicated concentrations of FITC-dextran (a) and nitazoxanide (b) was measured using UV-VIS spectrometry. Spectra were recorded from 370 to 750 nm and baseline-corrected using a distilled water reference. Peak absorbance values were recorded at 490 nm for FITC-dextran and 414 nm for nitazoxanide. The difference in absorbance between sample peaks and the absorbance of the ICM medium solvent (black line) is plotted as a function of concentration.

### 2.5. Automated sampling of the basolateral transwell chamber

A prototype automated sampling system (BioPhys-Concepts, Budapest, Hungary, biophys-concepts.com) configured to utilize four separate tubing systems: two delivered the fresh medium while the other two removed the medium from the basolateral and apical chambers. The dosing unit used two syringe pumps and a fluid multiplexer to measure and inject 1.2 mL cooled medium into the tube connected to the incubator unit. A peristaltic pump is used to remove the medium from the device. Samples were collected in 2 mL Eppendorf tubes in the cooled fraction collector of the sampling system. Within the culture incubator, the fresh medium was kept in a heat exchanger for 30 minutes before being injected into the culture well. The culture was placed on a rotary shaker to ensure homogeneity of the basolateral compartment. The rotary shaker induced an intermittent rotary motion at a radius of 10 mm and a top rotational speed of 120 rpm. The time delay between the complete removal of the basolateral medium and the injection of fresh medium was less than 10 sec. Between measurements, the tubing network was cleaned and disinfected overnight using an automated procedure. First, all fluid handling volumes were filled with 0.1% sodium hypochlorite for 20 minutes. After the hypochlorite was removed, the tubing network was rinsed six times with sterile distilled water.

### 2.6. UV-VIS Spectroscopy

VIS absorption spectra were recorded with a Shimadzu UV-2101PC double beam spectrophotometer (Shimadzu, Kyoto, Japan) in the 370-750 nm spectral range at a spectral resolution of 2 nm, using distilled water as reference. Samples were equilibrated for at least 30 minutes at normal atmospheric CO_2_ conditions and were loaded into 400 uL custom-fabricated cuvettes providing 1 cm optical path length. ICM was used in experiments involving spectroscopy analysis to avoid absorption from phenol red and serum proteins. DMEM culture media, as well as FITC and nitazoxanide, exhibit absorp-tion peaks below 550 nm (see Supplement Fig. 1). Thus, for baseline correction, a linear fit in the 550-750 nm spectral range was subtracted from the recorded spectra. Solvent-referenced spectra were obtained as the difference between the baseline-corrected spectra of the sample and the corresponding medium.

### 2.7. HPLC/MS

The amounts of chloroquine and petasol were determined using high-performance liquid chromatography (HPLC) coupled with high-resolution Orbitrap mass spectrometry (HR-MS). Analyses were performed on a Dionex Ultimate 3000 HPLC system (3000RS diode array detector (DAD), TCC-3000RS column thermostat, HPG-3400RS pump, SRD-3400 solvent rack degasser, and WPS-3000TRS autosampler) connected to an Orbitrap Q Exactive Focus mass spectrometer equipped with an electrospray ionization (ESI) source (Thermo Fisher Scientific, Waltham, MA, USA).

Chromatographic separation was achieved using a Kinetex C18 column (75 × 3 mm, 2.6 µm; Phenomenex, Torrance, CA, USA) at a flow rate of 0.3 mL/min and ambient column temperature. The injection volume ranged from 1.0 to 5.0 µL. Ionization was performed in positive ESI mode, and full MS scans were acquired at a resolution of 35,000 over an m/z range of 200–500. MS operational parameters were automatically optimized using the instrument software and were consistent with those described in our previous study [19].

Eluents were as follows: eluent A, 0.1% (v/v) formic acid in water; eluent B, acetonitrile containing 0.1% (v/v) formic acid. Two gradient programs were used:

Gradient 1: 0.0 min, 15% B; 3.0 min, 50% B (linear); 3.5 min, 90% B (linear).

Gradient 2: 0.0 min, 25% B; 7.0 min, 90% B (linear); 7.5 min, 90% B (isocratic). Chloroquine and petasol were analyzed using gradient programs 1 and 2, respectively, by detecting their protonated molecular ions. The differences between the measured and calculated mass-to-charge ratios (m/z) of the protonated species were within ± 1 ppm, confirming compound identity. For quantification, selected ion chromatograms (SICs) were generated by filtering at m/z 320 ± 0.5 for chloroquine and m/z 235 ± 0.5 for petasol. Peak areas in the SICs were manually integrated. Quantification was performed using an external calibration method. Chloroquine and petasol reference standards were obtained from Sigma-Aldrich (St. Louis, MO, USA) or isolated as described in our previous work (European Patent EP 4134075A1), respectively. Calibration curves were generated by linear regression within the concentration ranges of 0.0370–1.000 µM for chloroquine and 0.0370–4.750 µM for petasol. The amounts of chloroquine and petasol in the transwell chambers were then calculated using these calibration curves.

### 2.8. Sampling correction

Measured analyte amounts in sequential samples often deviate from the true transport profile due to surface adsorption–desorption effects and residual droplet retention, which together can cause cross-sample contamination [7,20]. To address these artifacts, we applied a linear recursive correction model, as detailed in [20]. The model parameters are obtained from cell-free calibration experiments, enabling the systematic removal of sampling-related distortions and yielding corrected concentration–time series suitable for downstream transport analysis.

### 2.9. Barrier models

For quantitative interpretation of corrected transport data, we employed compartmental models of increasing complexity [20]. The two-compartment model describes passive diffusion across the membrane by a single permeability parameter *P*_0_. Variables of this two compartment model are the concentrations in the apical and basal compartments, *c*_*A*_ and *c*_*B*_, respectively. The time evolution of these variables are given by:

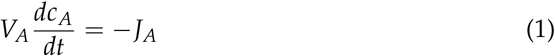

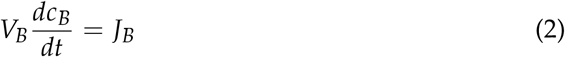

where *J*_*A*_ and *J*_*B*_ are the fluxes through the apical and basal surfaces, while *V*_*A*_ and *V*_*B*_ denote the volumes of the apical and basal compartments. Assuming simple diffusion through a thin, inert membrane

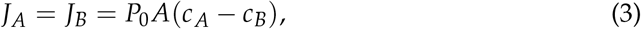

where *P*_0_ and *A* are the permeability and the area of the membrane.

To account for cellular processes beyond membrane passage, we extended this framework to a three-compartment model that incorporates an intermediate cellular space, parameterized by an effective layer height (*h*) governing reversible sequestration, an average diffusivity (*P*) and a metabolic loss rate (*m*) representing intracellular degradation. The model above are extended by a third, cellular, compartment in which the concentration of the free analyte is denoted by *c*_*C*_. We assume that the total amount of the analyte within cells, *M*_*C*_, is proportional to *c*_*C*_ as

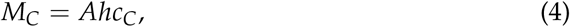

where *Ah* is an effective cell volume that accounts for reversible sequestration within the cell layer. The time development of the model is given by Eqs. (1), (2) and by

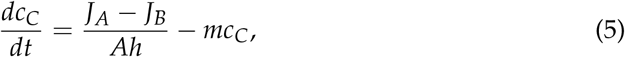

where parameter *m* characterizes an irreversible removal of the analyte within the cell layer through, e.g., metabolic degradation. The apical flux is determined by the cell membrane permeability, *P*, as

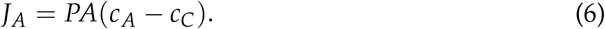

The flux at the basal cell surface is equal to *J*_*B*_, the flux through the porous membrane, yielding

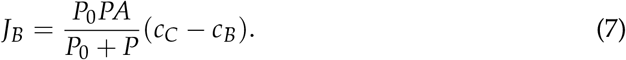

as described in detail in [20]. Parameters of both models were estimated by nonlinear regression against experimental sample sequences, allowing systematic comparison of passive and cell-influenced transport kinetics.

### 2.10. Computational methods

Python codes were used for processing UV-VIS spectra, fitting model parameters and for the analysis of sample sequences. As described in detail in [20], data analysis has been performed using the scipy python module version 1.6.0. The analysis scripts are available at https://github.com/aczirok/barrier-transport-tools.

## 3. Results

### 3.1. Barrier integrity of Vero E6 and Caco-2 monolayers

To assess the suitability of epithelial monolayers for barrier transport studies, transepithelial electrical resistance (TEER) was measured daily in both Vero E6 and Caco-2 cultures. The monolayers were grown on the apical surface of transwell inserts, and TEER measurements were performed using an automated system equipped with a chopstick electrode. Figure 2 summarizes the typical TEER values over time and compares the cultures used for transport measurements to similar cultures previously established in our laboratory.

**Figure 2.**
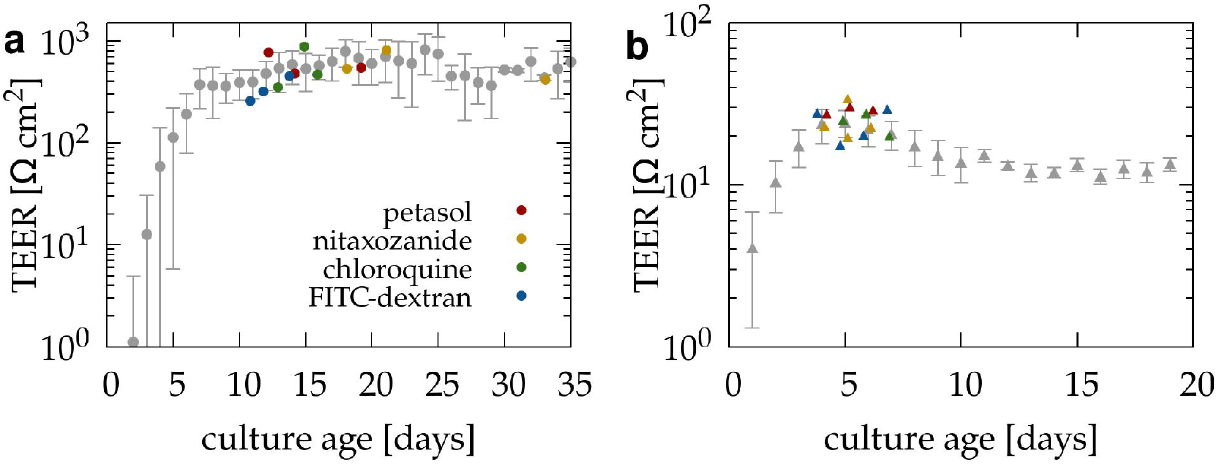
Maturation of epithelial barrier cell layers monitored by daily transepithelial electrical resistance (TEER) measurements. Gray symbols indicate mean TEER values and standard deviation obtained from n = 6 Caco-2 (a) and n = 17 Vero E6 (b) independent experiments, respectively, each with 3-6 cultures. After a lag phase marked by rising TEER values, cultures reached a stable plateau indicative of functional barrier formation, suitable for transport assays. Colored symbols mark the specific cultures selected for subsequent transport experiments.

Caco-2 cells exhibited a progressive increase in TEER during the first 7 days of culture, after which the resistance plateaued at values exceeding 400 O·cm^2^. Cultures maintained under these conditions remained stable for up to 30 days and were used for transport experiments. These TEER values are consistent with literature reports and reflect the formation of well-differentiated, polarized monolayers with intact tight junctions [21].

Vero E6 cultures, which characteristically form less resistive barriers, reached peak TEER values by approximately day 4 in vitro, exceeding 20 O·cm^2^. These values fall within the expected range for Vero E6 cells and indicate the presence of a continuous, confluent epithelial layer suitable for permeability studies. Thus, both cell models exhibited TEER profiles consistent with typical barrier-forming behavior, confirming their suitability for barrier transport assays.

### 3.2. measured barrier transport data

We evaluated analyte permeability across epithelial barriers for four compounds: petasol, chloroquine, nitazoxanide, and FITC-dextran (4 kDa). FITC-dextran is a commonly used negative control [22] that is not taken up by cells and can only cross the barrier via paracellular routes, making it a sensitive indicator of tight junction integrity. Nitazoxanide served as a positive control, as it is a clinically used antiviral compound known to cross epithelial barriers in vivo [23–25]. Chloroquine, another well-characterized compound [26,27], was included for comparison due to its lower permeability relative to nitazoxanide. Initial apical analyte concentrations were selected based on prior toxicology screens and detectability considerations, and were set to 25 µM for petasol, 12.5 µM for chloroquine, 25 µM for nitazoxanide, and 0.5 mg/mL for FITC-dextran. Individual transwell cultures were transferred into a fluidic system, which replaced the basolateral fluid compartment every 30 minutes and collected the effluent in a cooled sample collector to preserve analyte integrity. After two baseline samples were obtained, medium in the apical compartment was replaced with medium containing the analyte, and sampling continued for additional 6 hours. Analyte concentrations in the collected basolateral samples were quantified using either HPLC-MS or optical VIS spectroscopy, depending on the optical and chemical properties of the analyte.

Figure 3 shows the resulting time-dependent concentration profiles for the four compounds investigated. Each analyte was tested under four conditions: (i) across a Caco-2 monolayer, (ii) across a Vero E6 monolayer, (iii) across a cell-free transwell membrane to assess passive diffusion through the insert itself, and (iv) by direct injection into the culture medium without the presence of cells or a transwell insert. The latter served as a calibration procedure to evaluate potential cross-contamination in the sampling process – specifically, how much analyte may carry over from one sample to the next due to residual droplets or adsorption to tubing surfaces. As shown in Fig. 3, none of the compounds were detectable in the two baseline samples, confirming the absence of pre-existing contamination.

**Figure 3.**
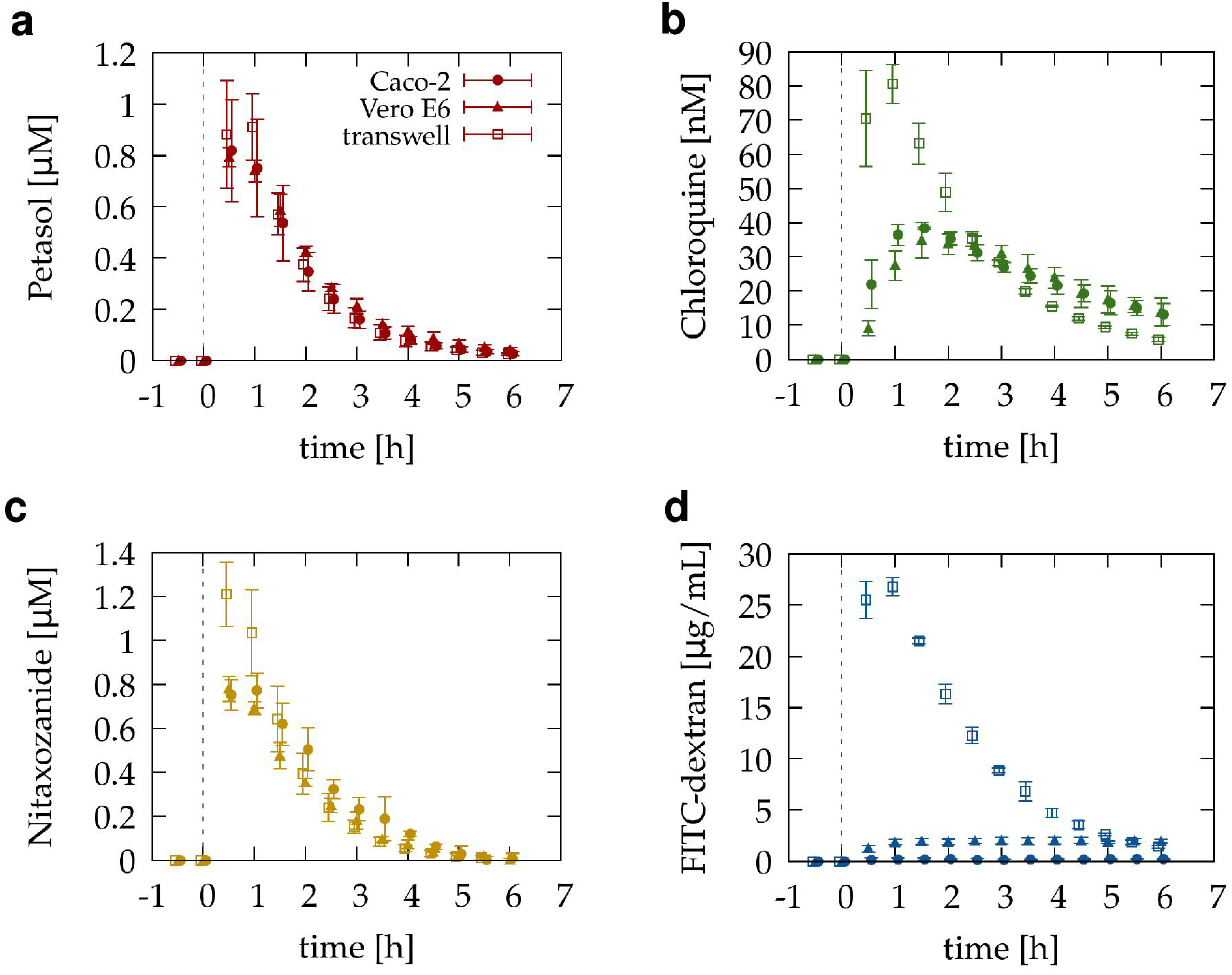
Time-resolved basolateral concentrations of four test compounds following apical administration to assess epithelial barrier transport. The translocation of antiviral compounds petasol (a), chloroquine (b), nitazoxanide (c) and the negative control FITC-dextran (d), was assessed through Caco-2 and Vero E6 monolayers, as well as cell-free transwell membranes. Test compounds were added to the apical compartment of Transwell inserts, and basolateral samples were collected and replaced every 30 minutes. Concentrations in collected samples were quantified using mass (a, b) and UV-VIS (c, d) spectrometry. Petasol showed similar transport across cell layers and cell-free membranes, whereas chloroquine and nitazoxanide exhibited delayed or reduced passage through the epithelial barriers. FITC-dextran was almost completely retained by the cell layers, confirming the integrity of the barriers.

For the cell-free membranes, all compounds followed a similar pattern: peak concentrations appeared in the third sample (immediately after the injection), followed by a gradual, exponential-like decline over time. FITC-dextran exhibited only minimal transport, with concentrations close to background levels – particularly across Caco-2 barriers, where it remained below the detection limit. Together with the TEER measurements, these data confirm that the epithelial barriers were intact and maintained functional tight junctions. Petasol and nitazoxanide displayed very similar concentration profiles across each experimental condition. Their transport was not notably affected by the presence of epithelial monolayers, indicating efficient permeability across both Caco-2 and Vero E6 cells. In contrast, chloroquine exhibited a markedly delayed transport in the presence of cell layers. The highest concentration was observed in sample 5, collected 90 minutes after apical administration, and the overall passage through the epithelial barriers was substantially slower compared to the cell-free condition.

The raw sample concentration data presented in Figure 3 were processed using a mathematical approach designed to estimate and correct for sample cross-contamination and analyte adsorption/desorption effects (see). The resulting total amounts of analyte recovered – combining the collected basolateral samples and the remaining apical medium at the end of the experiment – are shown in Figure 4. For each analyte, values were normalized to the average total amount detected for cell-free membranes to facilitate comparison across conditions. For petasol, the total amount detected remained unchanged irrespective of the presence of epithelial cell layers, suggesting minimal binding, degradation, or retention by the barriers. Nitazoxanide showed a moderate reduction in total amount across Vero E6 barriers, but not Caco-2 barriers, indicating potential cell-specific interactions. Importantly, neither petasol nor nitazoxanide accumulated in the apical compartment by the end of the transport experiments, consistent with efficient passage through the barriers. In contrast, chloroquine exhibited a pronounced reduction in the total amount recovered in the presence of cell layers, accompanied by substantial residual accumulation in the apical compartment. These observations point to slower transport, retention, or possible intracellular sequestration. Finally, as expected, most FITC-dextran remained in the apical compartment, reflecting its limited permeability; however, the presence of cells led to a detectable reduction in the total recovered amount, likely reflecting adsorption to cellular or membrane surfaces.

**Figure 4.**
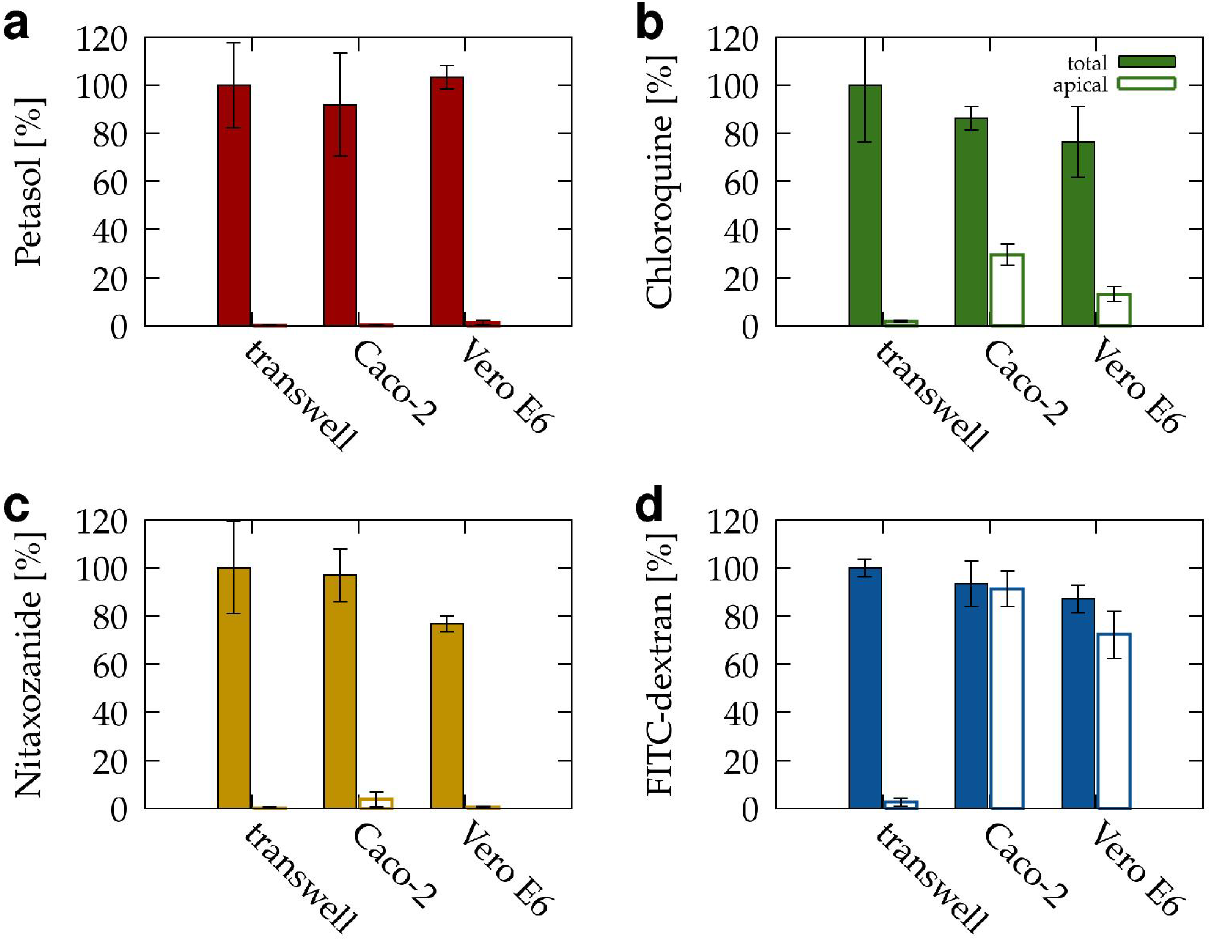
Recovery of test compounds after transport across Caco-2 and Vero E6 barriers. To assess whether compounds were reversibly or irreversibly affected (e.g., metabolized, bound, or retained) during transcellular transport, the total recovered drug in apical and basolateral samples was summed and normalized to the value obtained in cell-free transwells (filled bars). These values were compared with the drug recovered exclusively in the apical compartment (empty bars). Petasol was almost fully recovered in the basolateral fraction, indicating near-unhindered passage across both cell layers (a). Nitazoxanide also crossed the Caco-2 barrier efficiently, but its transport was moderately reduced by Vero E6 cells (c). In contrast, chloroquine transport was strongly impeded by both cell types, with substantial accumulation in the apical compartment (b). For the paracellular marker FITC-dextran, most recovery occurred in the apical compartment, confirming the barrier properties of both models (d). The apparent loss observed in the Vero E6 system likely reflects adsorption to the cell surface.

### 3.3. barrier transport fluxes

Based on the measured concentration of each compound in the apical compartment at the end of the transport experiment – and taking into account the total amount already transported and collected – we calculated the apical-to-basolateral concentration difference (Δc) for each time point, along with the corresponding amount of analyte transported across the barrier. As shown in Figure 5, in most systems, particularly in the case of cell-free membranes, the trans-barrier flux is proportional to c, consistent with passive diffusion. For epithelial barriers, this proportionality is primarily observed at later time points, when the concentration difference is lower and the system approaches steady-state conditions. A notable exception is chloroquine, which shows substantial deviations from this linear relationship, suggesting that its transport is influenced by additional factors beyond simple diffusion. FITC-dextran, on the other hand, exhibits negligible transport relative to the amount present in the apical compartment. Consequently, Δc remains effectively constant throughout the experiment, and the observed data scatter likely reflects measurement noise near the detection limit of optical spectroscopy.

**Figure 5.**
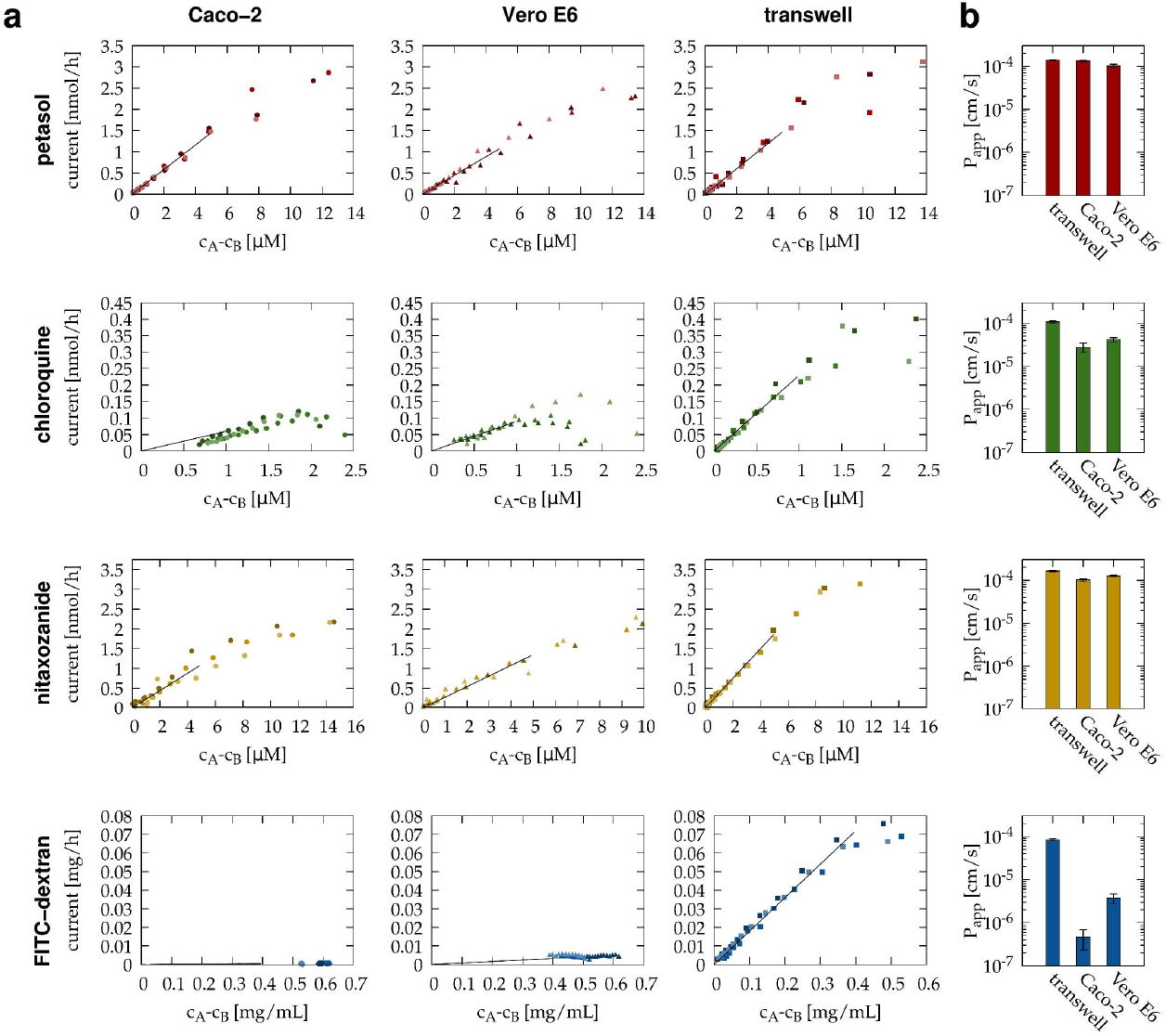
Characterization of compound transport across Caco-2 and Vero E6 barriers using a two-compartment model. Apical-to-basolateral concentration differences and corresponding fluxes were derived from compound amounts detected in basolateral samples after apical dosing. Data from three independent experiments are shown for each compound: petasol (red), nitazoxanide (yellow), chloroquine (green), and FITC-dextran (blue), across cell-free inserts (squares), Caco-2 monolayers (dots), and Vero E6 monolayers (triangles) (a). Linear flux–Δc relationships, observed for petasol and nitazoxanide, indicate passive diffusion. For chloroquine, deviations from linearity suggest additional processes such as adsorption, metabolism, or intracellular retention. FITC-dextran was largely retained by cell layers, yielding stable concentration differences throughout the experiment. In cases with moderate Δc, linear regression was applied to estimate apparent permeability coefficients (*P*_*app*_), which quantify transport efficiency (b).

The fitted proportionality line provides an estimate of *P*_*app*_, the apparent permeability of the barrier; its values are summarized in Figure 5d. Higher *P*_*app*_ values indicate more efficient transport across the barrier, while lower values reflect restricted permeability, typically due to tighter junctions or reduced analyte mobility.

### 3.4. Three compartment model parameters

The *P*_*app*_ value reflects the combined influence of several factors, including cell membrane permeability, analyte binding or retention, and transporter activity. It is based on the assumption that the trans-barrier flux is directly proportional to the concentration difference. However, the time-resolved sample series also permits the application of more detailed mathematical models that reveal more about the transport kinetics.

Here, we consider a three compartment model defined through Eqs. (1), (2), (5), (6) and (7). The fitting procedure yields four kinetic parameters: *P*_0_ and *P*_1_ are the permeabilities of the transwell membrane and the cell membrane, respectively. The effective cell layer height, *h*, accounts for the capacity of the epithelial layer to reversibly bind or transiently retain the analyte. Finally, the parameter *m* describes irreversible loss, such as through metabolic degradation.

As demonstrated in Fig. 6a, this simple model successfully captures some of the transient behavior observed in the time-resolved transport data and enables the estimation of cell- and analyte-specific kinetic parameters (Fig. 6b). The fitted values suggest that petasol readily traverses the cellular membranes, with minimal intracellular sequestration and no detectable metabolic degradation. In contrast, chloroquine displays reduced membrane permeability, and the fitted parameters indicate both significant intracellular binding and irreversible loss, consistent with metabolic degradation or sequestration.

**Figure 6.**
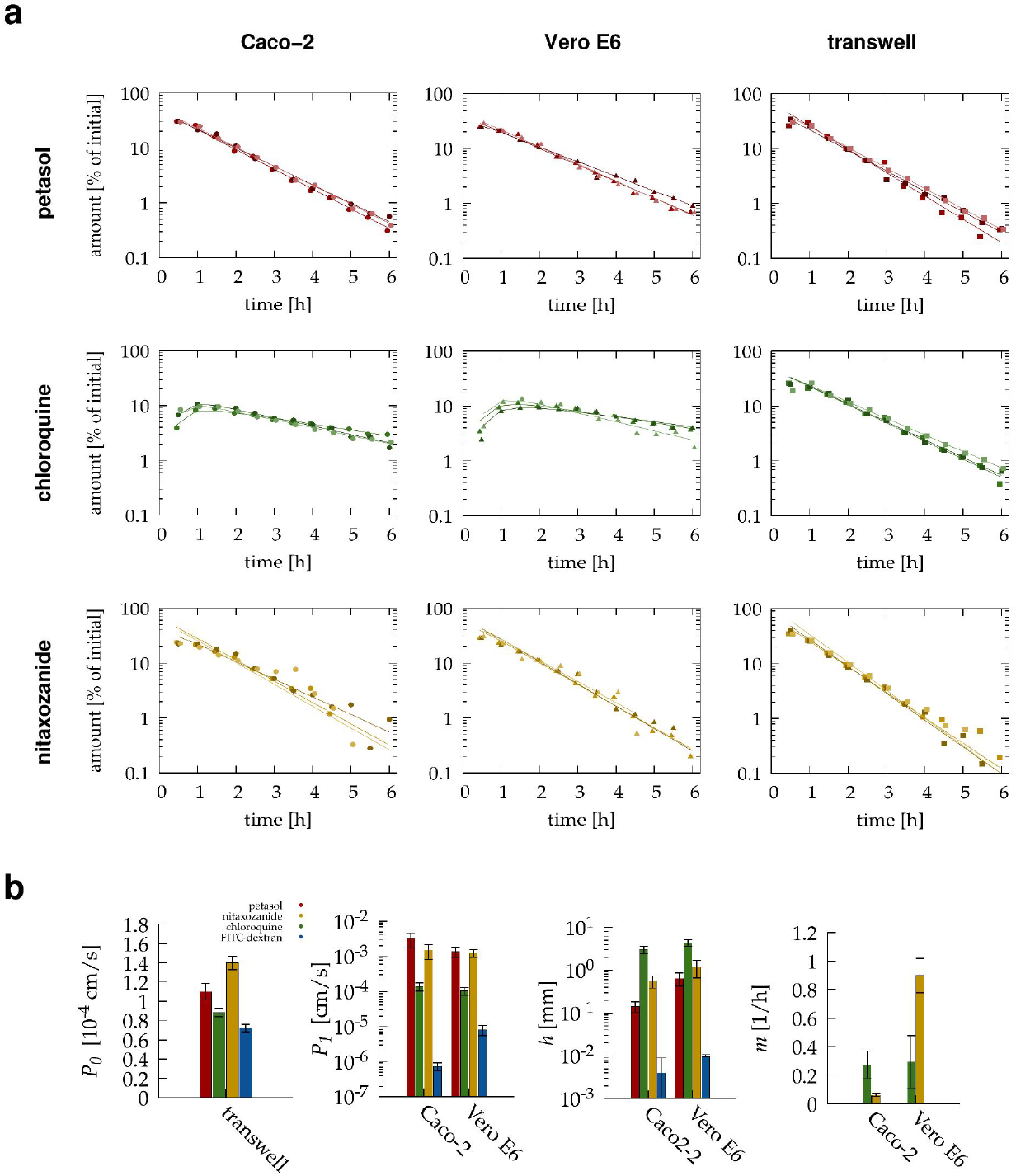
Three-compartment modeling of compound transport across Caco-2 and Vero E6 barriers. Basolateral recovery of petasol, nitazoxanide, chloroquine, and FITC-dextran was expressed as a percentage of the apical dose and fitted with a three-compartment transport model (a). The model parameters are: *P*_0_, Transwell membrane permeability; *P*_1_, cell membrane permeability; h, effective barrier height representing reversible processing that delays transport; and m, irreversible loss (e.g., metabolism). Parameter estimates with standard deviations from n = 3 independent experiments are shown for each compound–barrier combination (b).

## 4. Discussion

### 4.1. Petasol transport and comparison with reference compounds

A drug candidate’s ability to efficiently cross epithelial barriers is a prerequisite for systemic efficacy, particularly in orally administered therapies. Our findings indicate that petasol readily permeates both Caco-2 and Vero E6 monolayers, with transport profiles closely resembling or even better than those of nitazoxanide, a clinically used antiviral compound. These findings suggest that petasol can traverse biological barriers without significant hindrance from tight junctions, intracellular sequestration, or metabolic degradation. The high apparent (*P*_*app*_) and cell membrane (*P*_1_) permeability values highlight its favorable physicochemical properties for epithelial absorption. Efficient trans-epithelial transport supports oral bioavailability and facilitates tissue distribution in compartments protected by epithelial layers, such as the lungs or gastrointestinal tract—relevant sites for many viral infections.

Chloroquine, a well-characterized drug with reported epithelial transport properties, served as an internal control. Our results for chloroquine closely match previously published data for Caco-2 monolayers, including its moderate *P*_app_ value (2.1 · 10^*−*5^cm/s reported vs (2.8 ±0.7)· 10^*−*5^ measured) as well as the significant accumulation within epithelial cells [11]. This concordance supports the validity of our experimental approach and mathematical modeling framework.

In our experiments, FITC-dextran was a negative control, as it is not expected to cross intact epithelial barriers. Indeed, most dextran remained in the apical compartment, but a noticeable reduction in total recoverable amounts was observed in both Caco-2 and Vero E6 models (Fig. 4d). This apparent loss of dextran may reflect surface binding rather than translocation or degradation. Dextran has been reported to adhere to apical surface structures such as the glycocalyx, particularly in endothelial tissues [28]. Vero E6 kidney cells are known to express glycocalyx-like molecules [29,30], including glycosaminoglycans such as heparan sulfate, which have the potential to bind high molecular weight polysaccharides like dextran. Such binding could plausibly explain the incomplete recovery of FITC-dextran in our system, highlighting the need for caution when interpreting the loss of the total amount as cellular degradation.

### 4.2. Mechanistic insight from fitting time-resolved data

Our study goes beyond the estimation of *P*_app_ by analyzing time-resolved concentration profiles to extract additional kinetic parameters. These include the cell membrane permeability (*P*_1_), which encompasses both transcellular and paracellular routes, the effective cell layer height (*h*), which links intracellular analyte concentration to the total amount of analyte within the cell layer, and a metabolic degradation parameter (*m*), which captures irreversible loss during transport. Together, these parameters offer a more mechanistic understanding of how analytes interact with epithelial barriers and cellular environments.

However, the current three-compartment ODE model used to fit the data assumes instantaneous mixing and transfer across barriers, whereas in reality, spatial concentration gradients form within the cell layer and across the porous membrane. The gradual formation of concentration profiles can lead to a transient delay in transport, especially visible in the first sampling point [7]. This limitation of the model is responsible for the consistent overestimation of the first sample concentration seen in the fitted curves. Future work should thus consider spatially resolved diffusion models incorporating time-dependent concentration profiles within the epithelial layer and the membrane to better capture the early kinetics.

### 4.3. Implications for pharmacokinetics and therapeutic development

While our in vitro results are promising, further studies are needed to evaluate the pharmacokinetic behavior of petasol in vivo. These include assessments of absorption, distribution, metabolism, and excretion (ADME) using animal models, followed by pharmacodynamic correlation studies to determine effective tissue concentrations. Additional focus should be placed on metabolic stability and potential interactions with drug transporters, which may influence systemic availability or clearance rates.

Ultimately, the data presented here provide a critical first step in the preclinical characterization of petasol. Its efficient passage through epithelial barriers and the previously reported antiviral activity support its continued development as a therapeutic candidate. Integrating in vitro permeability data with in silico pharmacokinetic modeling will help refine dosing strategies and identify potential challenges before clinical testing.

## Data Availability Statement

All relevant data is included in the manuscript.

